# Accelerated mass transfer from frozen thin films during thin-film freeze-drying

**DOI:** 10.1101/2022.04.16.488553

**Authors:** Jie-Liang Wang, Manlei Kuang, Haiyue Xu, Robert O. Williams, Zhengrong Cui

## Abstract

Freeze-drying, or lyophilization, is widely used to produce pharmaceutical solids from temperature-sensitive materials but the process is time and energy inefficient. Herein, using *E. coli* as a model live organism, whose viability in dry powders is highly sensitive to the water content in the powders, we demonstrated that the drying rate of thin-film freeze-drying (TFFD) is significantly higher than that of the conventional shelf freeze-drying, likely because the large total surface area from the loosely stacked frozen thin films and the low thickness of the thin-films enable faster and more efficient mass transfer during freeze-drying. The highly porous nature and high specific surface area of the thin-film freeze-dried powders may have contributed to the faster mass transfer as well. Moreover, we demonstrated that TFFD can be applied to produce dry powders of *E. coli* and *L. acidophilus* with minimum bacterial viability loss (i.e., within one log reduction), and the *L. acidophilus* dry powder is suitable for intranasal delivery. It is concluded that TFFD technology is promising in addressing the time-and cost-inefficient issue of conventional shelf freeze-drying.

## 1. Introduction

Freeze-drying, or lyophilization, uses vacuums to remove solvents from frozen solutions or suspension. In the pharmaceutical industry, it is widely used to produce solids that are temperature-sensitive, such as biologics (Zhang et al., 2021). In a typical freeze-drying process, the bulk liquid is frozen in the primary container placed on the drying shelves, where the freezing time can range from minutes to hours, depending on the thermal properties of the container and the product. Next, the frozen sample is subjected to low pressure to remove bulk ice by sublimation, known as the primary drying phase. In the primary drying phase, it is assumed the mass transfer occurs only at the interface of ice and the dried portion of the sample, known as the “drying front”. The sublimation rate decreases over time as water molecules must travel longer through the porous channels (**Fig. 1A**). After removing the bulk ice, the shelf temperature is often ramped up to accelerate the desorption of bound water molecules to reach a residual water content of typically less than 3% for storage. This process is referred to as secondary drying, which is desorption-driven. The major downside of freeze-drying is that the drying process is time and energy inefficient as compared to other drying technologies such as spray drying and fluid bed drying. Because the mass transfer is limited to the air-sample interface, the large-scale freeze-drying process is particularly inefficient because the surface area to volume ratio (SA/Vol) decreases as volume increases in a fixed diameter container/vial, or remains the same if the filling height is fixed. Roughly, a freeze-drying batch on the liter scale may take several days to reach the target water content while spray drying would only take a few hours at most to complete. For this reason, freeze-drying is almost exclusively used for high-value, heat-liable materials with unsatisfactory stability in liquid formulations, such as inhalable insulin (Galderisi et al., 2020).

**Figure 1.**
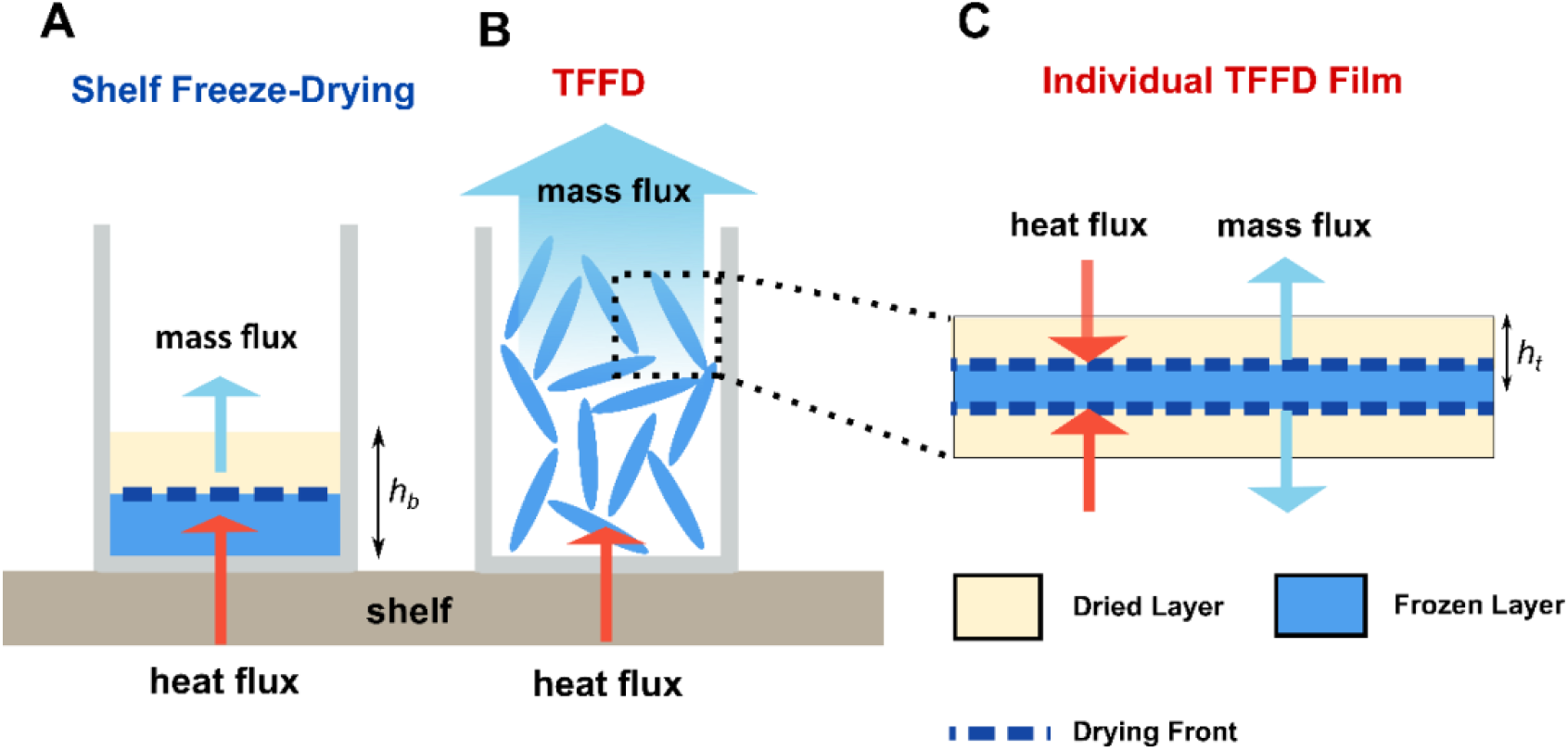
Proposed drying mechanism for thin-film freeze-drying (TFFD) in comparison to conventional shelf freeze-drying. **(A)** the heat and mass transfer of bulk a sample frozen on the shelf. **(B)** the heat and mass transfer of sample frozen by thin-film freezing (TFF) process. **(C)** the heat and mass transfer mechanism of an individual thin film prepared by TFFD.

To accelerate freeze-drying, various approaches such as drying cycle optimization with heat and mass transfer modeling (Bano et al., 2020; Kuu and Nail, 2009; Shivkumar et al., 2019), continuous freeze-drying (Capozzi et al., 2019), and spin freeze-drying (de Meyer et al., 2015) were introduced. Using heat and mass transfer modeling, the freeze-drying program is optimized by maximizing the shelf temperature to promote mass transfer without collapsing the lyophilization cake (Capozzi et al., 2019). In spin freeze-drying, the liquid-filled vials are spun along the longitudinal axis while freezing, so that the frozen sample uniformly coats the wall of the vials, increasing the SA/Vol ratio (de Meyer et al., 2015). Noticeably, spin freeze-drying can significantly shorten the total drying time (Lammens et al., 2021). However, the application of spin freeze-drying is limited to unit-dose vials and not suitable for bulk production.

TFFD is a two-step process with a thin-film freezing (TFF) step followed by drying inside a conventional freeze dryer (Johnston et al., 2006). During the TFF step, the liquid is dropped dropwise to a pre-cooled surface. Upon impact to the surface, the droplets spread and are rapidly frozen into thin films in milliseconds with cooling rates between 100-1000K. Next, the frozen films are filled into containers (e.g., glass vial, tray) (**Fig. 1B**) followed by water removal in a conventional freeze dryer. Thin-film freeze-dried powders (TFFD powders) are often brittle and highly porous, making them suitable for dry powder aerosolization (Wang et al., 2021). The TFF process and applications have been recently reviewed by Hufnagel et al (Hufnagel et al., 2022). Importantly, the total surface area of the frozen thin films is significantly larger than if the same volume of bulk liquid is frozen in a container/vial on the shelf of a freeze dryer. For example, for the bulk liquid in a vial frozen on the shelf of a freeze dryer, assuming that the frozen sample is cylindrical, then the volume and the total surface area are:

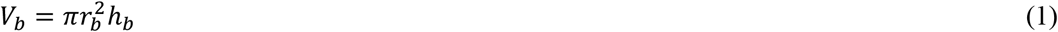

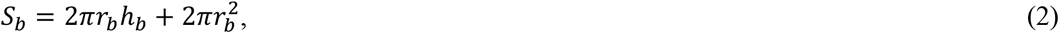

where *r* is the radius of the cylinder, and *h* is the height of the cylinder. Assuming the container surface has no water permeability, the mass transfer can only occur on or through the top of the frozen sample during freeze-drying (Muneeshwaran et al., 2022), then the effective mass transfer area is independent of the sample height:

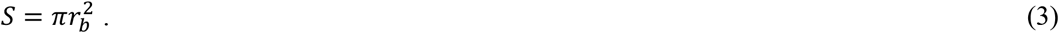

Therefore, if containers or vials with a fixed diameter are used, the effective surface area for mass transfer is fixed and the effective SA/Vol ratio can be expressed as:

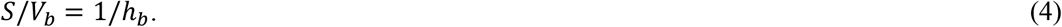

In practice, the freeze dryer is often designed with multiple shelves, so the height of the frozen liquid in the container is lower, often limited within 2 cm.

For comparison, in the TFF process, the bulk liquid is dispensed as small droplets and frozen into a large quantity of thin films. Assuming the frozen thin-films generated by the TFF process are cylindrical thin films or disks and the area intersections between individual cylinders are insignificant (Engstrom et al., 2008), then the total volume and surface area of the frozen thin-films are:

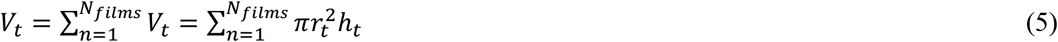

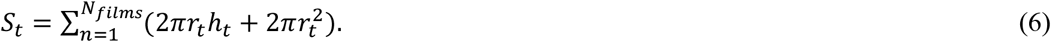

Therefore, the SA/Vol ratio would be much larger than the bulk frozen liquid in the container/vial when the number of the thin films is large, taking the advantage that mass transfer can occur on both sides of the films, and to a limited extent at the edge of the films (**Fig. 1C**). **Fig. 2** showed a theoretical plot of the SA/Vol ratio of samples processed by TFFD or shelf freeze-drying (FD). Clearly, the TFFD process would provide a higher SA/Vol ratio than shelf freeze-drying.

**Figure 2.**
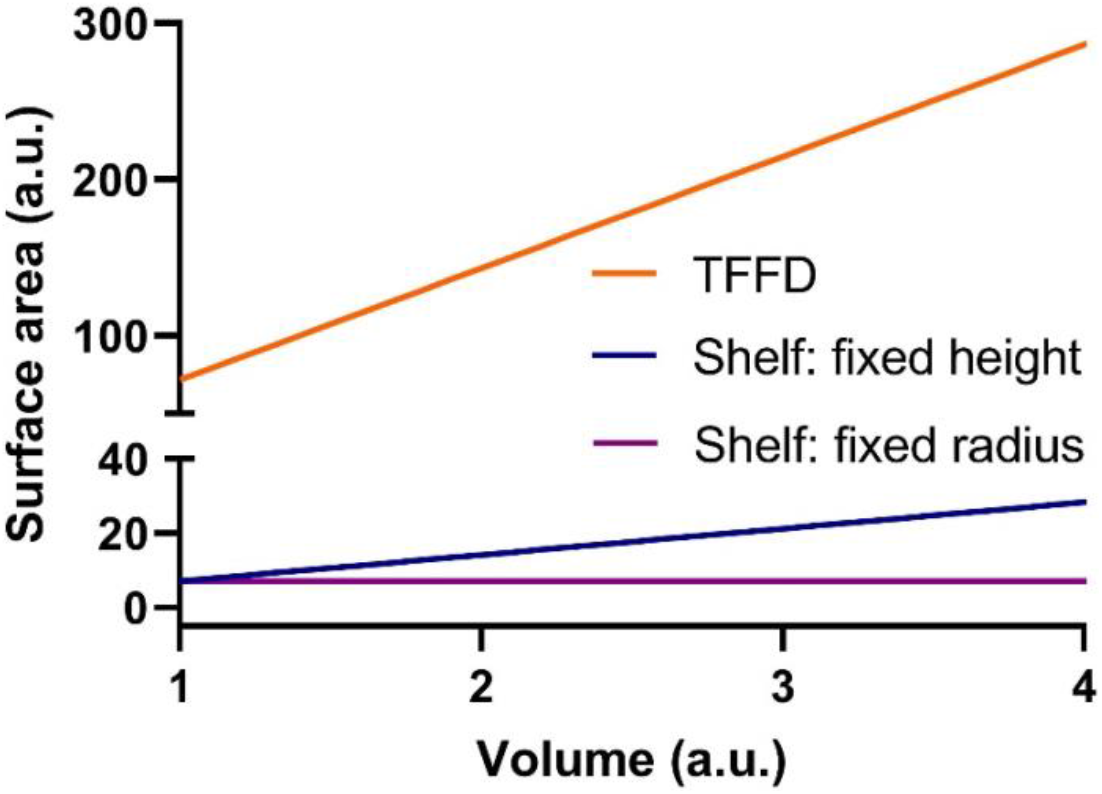
Theoretical relationships between the surface area for mass transfer to volume. **TFFD**: sample prepared by thin-film freezing (TFF) process. **Shelf, fixed height**: sample filled in a hypothetical vial and frozen on the shelf of a freeze dryer, increasing the sample volume by extending the vial radius while maintaining the sample height. **Shelf, fixed radius**: increasing the sample volume by increasing the sample height while the radius remains constant.

Besides the increased surface area, for individual frozen thin films, the heat and mass transfer model of drying slices from George and Datta also predicts the resistance to mass transfer in thin films is lower than in the bulk frozen samples because of the low thickness (George and Datta, 2002). In other words, the drying rate of individual frozen thin films should be high throughout the drying process due to its thin-film nature. Taken together, we hypothesized that the rate of mass transfer from frozen thin films produced by TFF is faster and more efficient than from bulk frozen samples generated by shelf freezing. TFF can overcome the limitation of current freeze-drying process acceleration techniques without the constraints of container shape.

Lastly, the TFFD process and shelf freeze-drying have different heat transfer mechanisms. In shelf freeze-drying, the shelf itself supplies heat to the frozen sample via the bottom of the container/vial to promote mass transfer, and the heat transfer is mainly by conduction, and to a less extent by convection and radiation. However, when freeze-drying the frozen thin films that only intercept one another or the container/vial at the edge, and with large air space among frozen thin films. Radiation and convection, instead of conduction, should play a more important role in heat transfer. Because ice sublimation and desorption process are promoted by heat (Pikal et al., 1990), the reduced conduction may decelerate the drying process. Moreover, the rapid freezing in the TFF process produces small ice crystals, which form narrower water channels that might have lower water diffusivity. This is different from bulk freezing, where larger ice crystals were produced by slow freezing and sometimes encouraged by annealing (Nakagawa et al., 2018).

To test our hypothesis, we performed freeze-drying experiments comparing the shelf freeze-drying and TFFD using a formulation containing live bacteria. We selected bacterial viability as the critical quality attribute (CQAs) because the water content is particularly important for live cell viability, and low water content can irreversibly damage cells (Tunnacliffe et al., 2001). Therefore, the water content for live-cell formulations should be tightly controlled to balance the viability and shelf-life. In comparison, small molecule formulations often use prolonged drying programs to minimize the water content, as high-water contents in the formulation may cause poor flowability, hydrate formation, and recrystallization (Zografi, 1988). Controlling the water content increases the process robustness but it is less significant in terms of CQA compared to live-cell formulations. Selecting the appropriate water content depends on the desiccation tolerance of bacteria. In general, bacillus has better desiccation tolerance than *E. coli* (Cruz Barrera et al., 2020). The tolerance can also be improved by optimizing the cryoprotectant formulation (Louis et al., 1994), or by pre-conditioning bacteria in high osmotic environments prior to freeze-drying (Manzanera et al., 2004). Therefore, a systematic study was first designed to identify critical process parameters (CPPs) that contribute to the *E*.*coli* viability in the TFFD process, which included (1) pre-treatment of the bacteria with a high salt broth before freezing, (2) thin-film freezing temperature, and (3) secondary drying time. Then we used the optimized CPPs to test if water removal from the bacterium-containing frozen thin-films prepared by TFF is faster and more efficient than from the bulk frozen bacterial suspension generated by shelf freezing while maintaining the desiccation-sensitive *E. coli* viability. Finally, we also tested the feasibility of applying the TFFD technology to prepare a dry powder formulation of *L. acidophilus*, a Gram-positive bacterium, for potential intranasal delivery (Szatraj et al., 2017), while minimizing bacterial viability loss.

## 2. Materials and Methods

### 2.1. Materials

Luria-Bertani (LB) broth and sodium chloride were from Fisher Bioreagents (Fair Lawn, NJ). De Man, Rogosa, and Sharpe (MRS) broth was from Oxoid Ltd (Basingstoke, Hampshire, UK). M9 broth, calcium chloride, magnesium chloride, and L-leucine were from Sigma-Aldrich (St Louis, MO). Sucrose was from Merck KGaA (Darmstadt, Germany) and sodium ascorbate was from Spectrum Chemical Mfg. Corp. (Gardena, CA). *E. coli* DH5α was from Thermo Fisher Scientific (Waltham, MA). The pUC19 plasmid was transformed into *E. coli* DH5α for ampicillin resistance. *Lactobacillus acidophilus* (Moro) Hansen and Mocquot (#4356) were from the American Type Culture Collection (Manassas, VA).

### 2.2. Bacterial viability and formulation

To test bacterial viability, the lyophilized bacterial powder was reconstituted to the pre-lyophilized volume with water with gently shaking, and then the viable bacteria were counted by the standard serial dilution method using either LB plate for *E. coli* or MRS plate for *L. acidophilus*. The *E. coli* formulation for freeze-drying was selected based on literature with modifications (Wessman et al., 2013). The formulation contains 10% w/v sucrose, 0.34% w/v M9 minimal salt supplied with CaCl_2_ and MgSO_4_, and 0.075% w/v leucine. M9 minimal salt was included to simulate the carry-over of growth media during the harvesting step during large-scale production. Leucine was included to reduce the hygroscopicity of dry powders (Li et al., 2016).

### 2.3. Preparation of *E. coli* frozen thin-films

*E. coli* was inoculated to LB broth containing 50 mg/mL ampicillin and incubated overnight at 37 °C with agitation and then diluted with additional LB broth containing ampicillin. The culture was incubated at 37 °C with agitation until the OD reached 0.8. Next, the bacterial culture was harvested by centrifugation at 6300 rcf for 7 min. The pellet was resuspended either in high osmotic media (M9 broth supplemented with 0.4 M of NaCl) for the osmotic shot, or directly in the cryoprotectant formulation. For bacterial culture underwent osmotic shot, the bacteria were incubated for an additional 2 h at 30 °C before they were harvested and resuspended in the cryoprotectant formulation. The resuspended bacteria were held on ice and used within 6 h. For the TFF process, the fresh bacterial suspension was frozen dropwise on a rolling stainless-steel drum precooled to different temperatures. The volume of the individual liquid drop was estimated by dividing the liquid volume by the number of drops dispensed. With a 21-gauge syringe, 1 mL of the liquid formulation was dispensed with approximately 78 drops (n=3). For each mL of liquid, the frozen films were collected into a 5 mL borosilicate glass serum vial and stored in a -80 °C freezer.

Detailed experimental conditions are in **Table 1**. In the freeze-and-thaw experiment, the frozen thin films in vials were held in a -80 °C freezer for 2 h and then thawed in a 37 °C water bath with gentle shaking.

**Table 1.** Summary of process parameters used in this study

| Condition | Bacteria | Freezing Method | Osmotic Shock | TFF Temperature | Drying Cycle |
| --- | --- | --- | --- | --- | --- |
| 1 | *E.coli* | TFF | Yes | -40 °C | 2 |
| 2 | *E.coli* | TFF | No | -40 °C | 2 |
| 3 | *E.coli* | TFF | Yes | -80 °C | 2 |
| 4 | *E.coli* | TFF | Yes | -140 °C | 2 |
| 5 | *E.coli* | TFF | Yes | -40 °C | 3 |
| 6 | *E.coli* | TFF | Yes | -40 °C | 4 |
| 7 | *E.coli* | Shelf | Yes | NA | 2 |
| 8 | *E.coli* | Shelf | Yes | NA | 3 |
| 9 | *E.coli* | Shelf | Yes | NA | 4 |
| 10 | *E.coli* | Shelf | Yes | NA | 1 |
| 11 | *E.coli* | TFF | Yes | -40 °C | 1 |
| 12 | *E.coli* | TFF | Yes | -80 °C | 1 |
| 13 | *E.coli* | TFF | Yes | -140 °C | 1 |
| 14 | *L. acidophilus* | TFF | No | -40 °C | 5 |

### 2.4. Freeze-drying cycles and film size measurement

The drying cycles used in this study are shown in **Table 2** with pressures set to 100 mTorr. The vials containing frozen bacterial films were transferred into a VirTis AdVantage Pro Freeze Dryer (SP Industries, Warminster, PA). For TFF samples, the freeze dryer shelf was precooled to -30 °C before sample placement. For shelf freeze-drying samples, the bacteria suspension was frozen in the freeze dryer. After the drying cycle, the diameter of dried thin films was measured using a caliper from a random selection of 10 intact films.

**Table 2.** Freeze-drying cycles used in this study.

| Step | Cycle 1 | Cycle 2 | Cycle 3 | Cycle 4 | Cycle 5 |
| --- | --- | --- | --- | --- | --- |
| 1 | Hold at -30 °C for 960 min. |  |  |  |  |
| 2 | End | Ramp to 25 °C at 1 °C/min |  |  | Ramp to 4 °C at 1 °C/min |
| 3 |  | End | Hold at 25 °C for 60 min | Hold at 25 °C for 300 min | Hold at 4 °C for 60 min |
| 4 |  |  | End | End | End |

### 2.5. Preparation of *L. acidophilus* dry powder

The model for lactic acid bacteria (LAB) used in this study was *L. acidophilus. L. acidophilus* bacteria were inoculated in 4 mL MRS broth and cultured in an anaerobic incubator at 37 °C for 48 h. Then the culture was transferred to 40 mL MRS broth and cultured for an additional 48 h. The bacteria were harvested and resuspended in a sterile solution containing 200 g/L sucrose, 9 g/L sodium chloride, and 5 g/L sodium ascorbate (Fonseca et al., 2015). The suspension was frozen with the TFF process temperature of -40 °C, followed by solvent removal using freeze-drying cycle 5 (**Table 2**).

### 2.6. X-ray powder diffraction (XRD), water content, and specific surface area (SSA) measurements

XRD was conducted using a Rigaku Miniflex 600 X-ray diffractor (Rigaku, Woodlands, TX, USA), using a monochromated Cu K radiation source (λ = 1.54056 Å) with an accelerating voltage of 40 kV at 15 mA. Samples were scanned in a continuous mode with a step size of 0.04° over a 2θ range of 3–60° at a scan speed of 2°/min, and a dwell time of 2 s. The water content of the lyophilized powder was measured using a Mettler Toledo C20 Coulometric KF-titrator (Columbus, OH). The SSA of the powder was measured using the Brunauer-Emmett-Teller (BET) method with a Quantachrome AutoFlow BET+ surface area analyzer (Anton Paar, Graz, Austria). To minimize the interferences of moisture on BET measurement, the powder sample was left to dry for an additional 24 h (100 mTorr, 25 °C shelf temperature) after the normal freeze-drying cycle.

### 2.7. Microscopy and aerosol particle size analysis

The freeze-dried sample was loaded on a carbon tape and then coated with 60:40 Pd/Pt using an EMS 500X Sputter Coater (Electron Microscopy Sciences, Hatfield, PA) and then examined using a Hitachi S-5500 scanning electron microscopy (Tokyo, Japan). For particle size analysis, the TFFD powder with *L. acidophilus* was passed through a test sieve with 300 µm openings and then loaded into a DeVilbiss Model 119 Powder Blower (DeVilbiss Healthcare, Port Washington, NY). The geometric diameter of the aerosols was measured by blowing the powder into a Spraytec laser diffraction system (Malvern Panalytical, Worcestershire, UK).

### 2.8. Statistical Analysis

For comparing the means of two groups, Student’s t-test was used. For bacterial viability involving multiple comparisons with three or more groups, the Welch and Brown-Forsythe ANOVA was performed, followed by Dunnett’s T3 post hoc multiple comparison test. Square root transformation was performed when the residual plot and Q-Q plot indicated poor-fitting with raw data. For multiple comparisons other than viability, an ordinary ANOVA test was used, followed by the Bonferroni post hoc multiple comparison test. In this study, the null hypothesis was rejected at p<0.05, denoted by *.

## 3. Results and discussion

### 3.1. Effect of pre-treatment in a high osmolarity medium, thin-film freezing temperature, and secondary drying time on the viability of *E. coli* after being subjected to TFFD

Considering the TFF process is a continuous process, we compared the bacterial viability when the *E. coli* suspended in the cryoprotectant formulation were held on ice for 0 h vs. 6 h and found that there was no significant difference in the bacterial viability (P > 0.05, n = 3). Therefore, the harvested bacteria used in the present study were frozen within 6 h. The log reduction of the *E. coli* viability was around one after the bacteria were subjected to the following TFFD protocol, i.e., resuspending in the cryoprotectant formulation, thin-film freezing at -40°C and then freeze-drying using the cycle 2 (i.e., -30 °C for 16 h, ramping up to 25 °C at 1 °C/min) (**Fig. 3A**). Incubating the bacteria in a high osmotic environment before subjecting them to TFFD (see **Table 1**, #1 vs. 2) helped to decrease the viability reduction by approximately 0.5 log CFU (p < 0.05, **Fig. 3A**). This is consistent with the literature that bacteria can accumulate osmolytes in high osmotic environments, improving their tolerance to subsequent freezing and drying stresses (Cruz Barrera et al., 2020; Garcia de Castro et al., 2000; Tunnacliffe et al., 2001).

**Figure 3.**
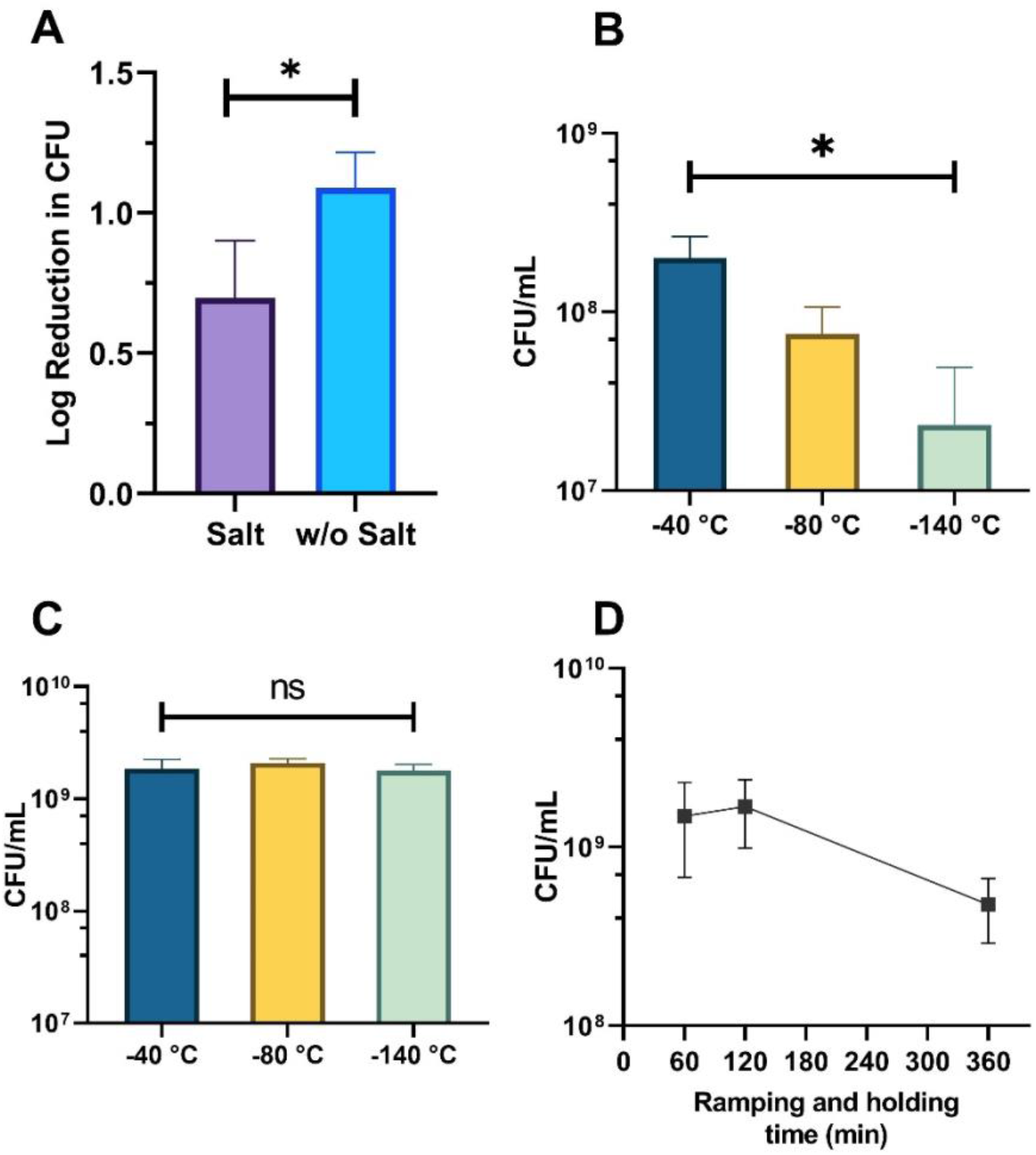
Identification of critical process parameters (CPPs) affecting the *E. Coli* viability in formulation processed by TFFD. **(A)** bacterial viabilities after the TFFD process, with or without incubating *E. coli* in M9 broth with 0.4 M sodium chloride for 2 h before being subjected to TFFD (n = 6, \**P* < 0.05). **(B)** Effects of -40 °C, -80 °C, and -140 °C TFF processing temperatures on bacterial viability after being subjected to TFFD (n = 6, \**P* < 0.05). **(C)** Bacterial viability after thin-film freeze-and-thaw without vacuum drying, using -40 °C, -80 °C, or -140 °C TFF processing temperatures. **(D)** Effects of secondary drying time on bacterial viability using -40 °C TFF processing temperature. After 16 h of primary drying at -30 °C, shelf temperature was ramped to 25 °C at 1 °C/min, followed by holding at 25 °C for an additional 0 min, 60 min, or 300 min (n = 3).

A key advantage of TFFD is that the cooling rate can be tailored by controlling the surface temperature of the precooled surface. We then studied the effect of processing temperature, and thus the freezing rate, on the viability of the *E. coli* after being subjected to TFFD (**Table 1**, #1, 3, 4, 7). As shown in **Fig. 3B**, the viability of the bacteria was retained best with a processing temperature of -40 °C, while the viability loss increased when the freezing temperature was reduced to -80 °C and -140 °C. To test whether the loss of bacterial viability occurred at the freezing or the subsequent drying step, we evaluated the viability of the *E. coli* in a freeze-and-thaw experiment, where bacteria samples were subjected to thin-film freezing at - 40°C, -80°C, or -140°C and then thawed without drying. Interesting, there was no significant difference in the bacterial viability between the processing temperature groups (**Fig. 3C**), clearly indicating that the processing temperature affected the viability of the bacteria in the drying step rather than the freezing step.

In conventional shelf FD, secondary drying mostly helps remove the residual bound water via desorption to promote stability during long-term storage and prevent cake collapse. However, *E. coli* is particularly sensitive to desiccation, making the selection of drying endpoints critical (Tunnacliffe et al., 2001). We examined the effect of secondary drying time on *E. coli* viability from samples prepared by thin-film freezing by extending the secondary drying time from 0 h to 1 h and 5 h using the “sample thief” mechanism (i.e., after the shelf temperature was ramped to 25 °C, removing vials from the freeze-dry after holding for 0, 1 or 5 h at 25 °C, **Table 1**, #5, 6). As shown in **Fig. 3D**, the viability of the bacteria was reduced significantly as the secondary drying time was extended to 5 h (**Table 1**, #1, 5, 6). Based on **Fig. 3D**, a secondary drying time at 25 °C for 0-1 h was appropriate for the thin-film frozen *E. coli* bacteria. Since the tolerance of desiccation depends on the strain of bacteria and the pre-treatments to the bacteria, there is not a general rule on selecting the appropriate water content. For *E. coli*, a water content of approximately 1.5% was reported to provide good stability and viability (Garcia de Castro et al., 2000). The requirement of maintaining the water content is unique to drying live cells where the bound water molecules are essential to the integrity and viability of the cells.

### 3.2. Water is removed from frozen thin films faster and more efficiently

Suspecting that the 5 h secondary drying time was too long so that the water content in the resultant dry powder was too low for *E. coli* to maintain their viability (**Fig. 3D**), we determined the water content in *E. coli* bacterial powders prepared by TFFD or conventional shelf FD using the “sample thief” mechanism by removing vials from the freeze-dryer immediately after the temperature is ramped to 25°C, or after 1 or 5 h of secondary drying at 25 °C (**Table 1**, #1, 5-9). As shown in **Fig. 4**, the water content in the TFFD powder was 3.4% after 16 h of primary dying and ramped to 25 °C at 1 °C/min. The water content was further reduced to 0.62% after 60 min of the secondary drying at 25 °C, and even further to 0.37% after 5 h of the secondary drying at 25 °C. Taken the data in **Fig. 3D** and **Fig. 4** together, it is likely that the water content in the TFFD powders must be above 0.5% to minimize bacterial viability loss after TFFD; reducing it to below 0.5% led to more viability reduction. In contrast, water removal from the shelf frozen samples was significantly slower and less efficient. For example, the water content in shelf FD powder remained more than 2% even after 5 h of secondary drying (**Fig. 4**), clearly demonstrating that compared to shelf FD, the TFFD process was more efficient in removing water during the drying step. Note that the water content in the TFFD powder was already below 4% at the first time point (i.e., with 0 h of secondary drying) while the water content in the shelf FD sample was still over 8% (**Fig. 4**). This suggested the desorption-driven secondary drying mechanism took place much earlier in TFFD than in shelf FD.

**Figure 4.**
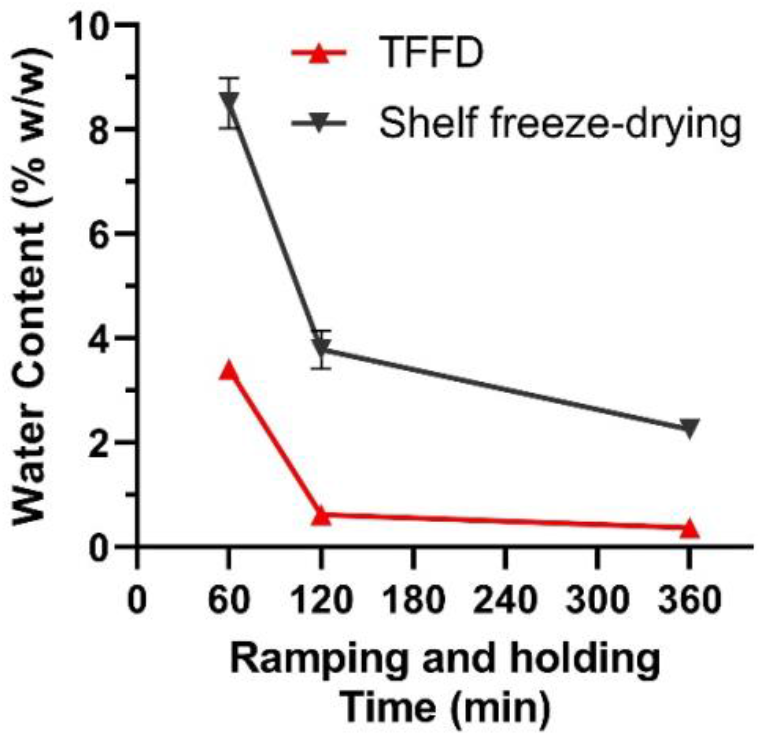
Water contents for samples processed by shelf freeze-drying or TFFD. Samples were subjected to 16 h primary drying at -30 °C, followed by ramping to 25 °C at 1 °C/min, and held for 60 min, 120 min, or 360 min at 25 °C (n = 3, 100 mTorr).

We then calculated the surface area to volume ratio of the thin films produced by TFFD. By dividing the number of droplets over the total volume, the volume of each liquid droplet was estimated to be 12.8 µL. The mean diameter of over 10 TFFD films processed at -40 °C was measured to be 7.34 ± 0.41 mm. The film thickness was 0.303 mm calculated using equation (1) and the surface area of individual TFF film was approximately 0.916 cm^2^ as calculated using equation (2). Therefore, the total surface area of 1 mL of frozen liquid is approximately 71.91 cm^2^ using equation 6. In comparison, for 1 mL of the same bacterial in liquid suspension frozen in a 5 mL serum vial with an inner diameter of 20 mm, the area of the surface of the frozen suspension that faces the air is approximately 7 cm^2^. Therefore, if the frozen thin films from 1 mL of bacterial suspension do not break and do not tightly stack on one another when randomly placed in a vial, then the total solid-air surface area of the frozen thin films was about 10 times larger than the solid-air surface area of the 1 mL shelf frozen suspension in a 20 mm vial. In reality, the total surface area of the thin films is expected to be smaller than the summation of all cylindrical thin films/disks due to their contact with the inner wall of the container and with other films and the stacking of films. Nonetheless, the total solid-air surface area of the frozen thin films should still be significantly larger than the shelf frozen bacterial suspension.

### 3.3. The thin-film nature of the frozen thin-films is likely responsible for the efficient removal of water during freeze-drying

To study the drying mechanism during the primary drying phase exclusively, we thin-film froze the bacteria at -40 °C, -80 °C, or -140 °C processing temperature and dried the frozen thin-films at -30 °C, 100 mTorr for 16 h (i.e., primary drying only, **Table 1**, #10-13) and measured water contents and bacterial viability, along with bacteria that were subjected to shelf FD as the control. The resultant shelf FD samples partially collapsed after being removed from the freeze dryer, indicating higher water content. Results shown in **Fig. 5A** again confirmed that compared to shelf FD, the TFFD process was more efficient in removing water from the frozen samples. Interestingly, the water content in the TFFD powders thin-film frozen at -80 °C or -140 °C were significantly lower than that in the TFFD powders thin-film frozen at -40 °C (**Fig. 5A**), indicating that certain properties other than the total air-solid surface area enabled the thin-films processed at -80 °C or -140 °C to be dried faster than the frozen thin-films processed at -40 °C. The lower water content also impacted the bacteria viability. Data in **Fig. 5B** showed that low thin-film freezing temperatures (−80 °C or -140 °C) led to lower *E. coli* viabilities in the TFFD powders frozen at -40 °C processing temperature.

**Figure 5.**
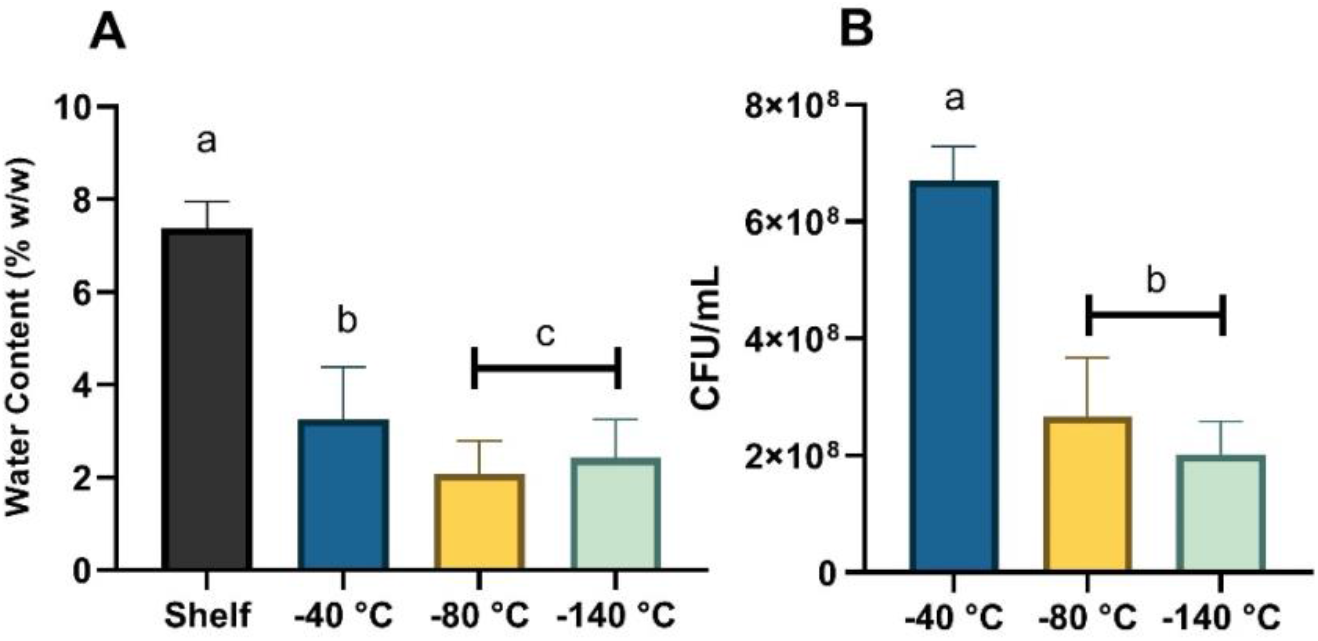
Comparison of water content and bacterial viability in dry powders prepared with different TFF processing temperatures. **(A)** Effects of TFF processing temperatures on water content after 16 h primary drying at -30 °C shelf temperature, compared to shelf freeze-dried samples (n = 3, ^a, *b, c*^*P* < 0.05). **(B)** Effects of TFF processing temperatures on *E. coli* viability at -30 °C shelf temperature for 16 h (n = 3, ^a, *b*^*P* < 0.05).

We then investigated certain microscopic physical properties of the resultant powders. We first examined the microstructures of the powders prepared by TFFD at different TFF processing temperatures or by shelf FD by scanning electron microscopy (SEM). Overall, it appears that TFFD powders, especially the one processed at -140 °C, were more porous than the shelf FD powder (**Fig. 6**). This is consistent with previous findings with TFFD tacrolimus dry powders (Sahakijpijarn et al., 2020). To further confirm the high porosity in the TFFD powders, the SSA values of the powders were determined. As shown in **Fig. 7A**, shelf FD powder had a much lower SSA value than the TFFD powders. Among the three different TFFD powders, the SSA value of the powder prepared at -40 °C was significantly lower than that prepared at -80 °C and -140 °C, while there was not a significant difference in the SSA values between the TFFD powders prepared at -80 °C and -140 °C (**Fig. 7A**). Finally, the XRD diffractograms of the TFFD powders prepared at all three freezing temperatures were similar (**Fig. 7B**) and consistent with the amorphous sucrose as previously reported (Nunes et al., 2005), indicating that the solid-state of the excipients in the TFFD powders was not likely related to the bacterial viability and drying efficiency.

**Figure 6.**
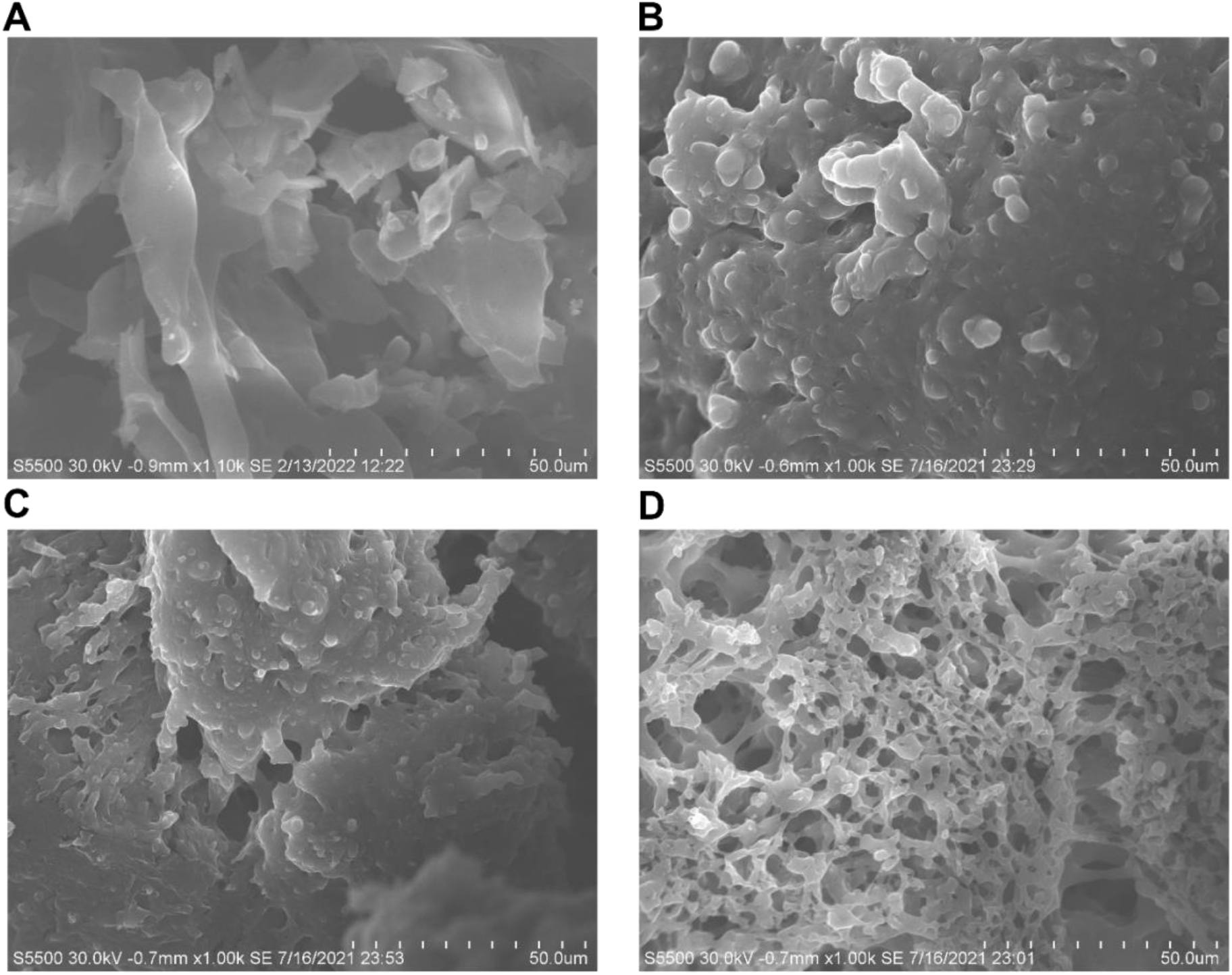
SEM images of freeze-dried *E. coli* powders processed by different freezing methods. **(A)** shelf freeze-dried samples frozen with a cooling rate of 0.5 °C/min and held at -30 °C for 60 min before drying. **(B-D)**, TFFD samples frozen with TFF processing temperatures of -40 °C, -80 °C, and -140 °C, respectively.

**Figure 7.**
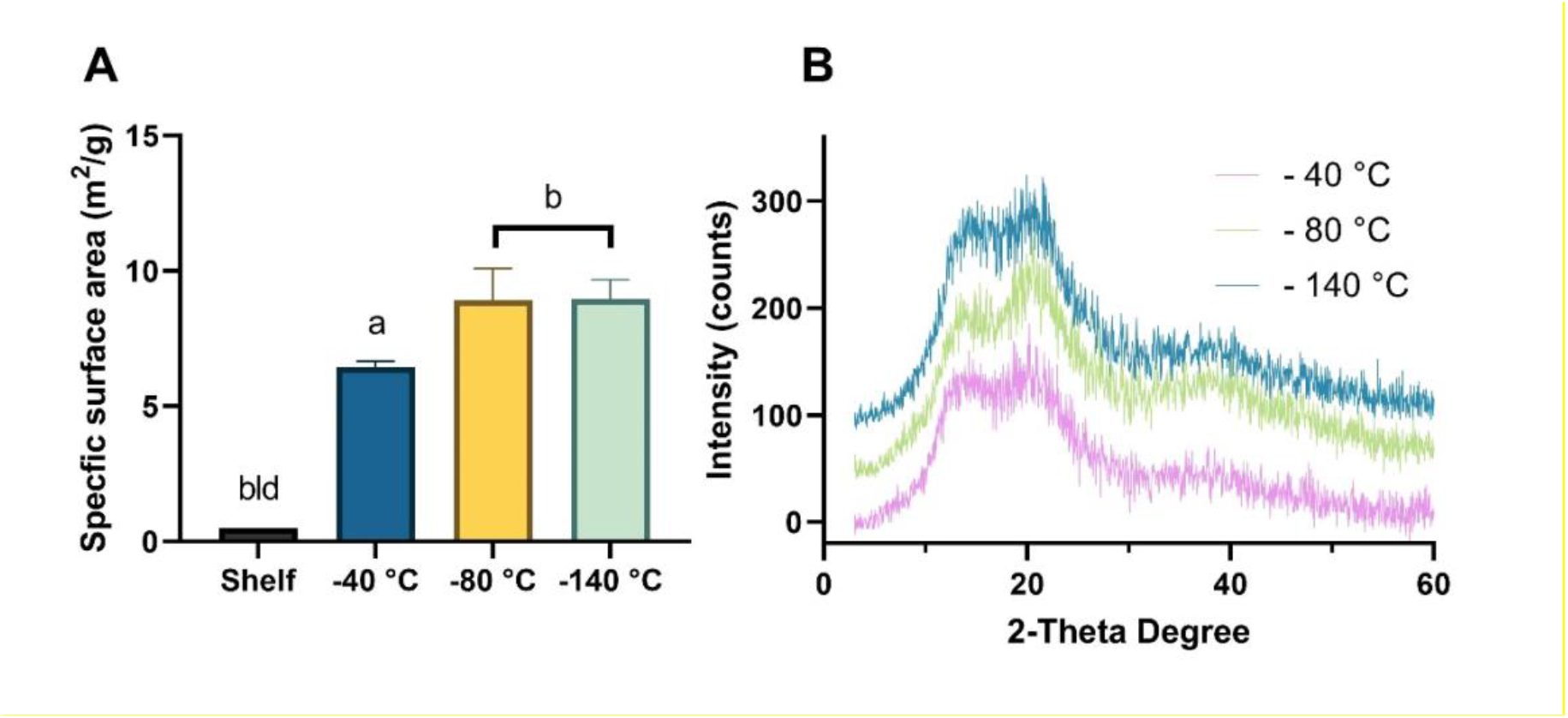
Comparing physical properties of powders prepared at -40 °C, -80 °C, or -140 °C TFF processing temperatures. **(A)** Effects of TFF processing temperatures on powder’s specific surface area, compared to shelf freeze-dried samples (n = 3, ^a, b^P < 0.05, bld: below the detection limit and excluded from statistical analysis). **(B)** XRD chromatograms of *E. coli* powders processed at different TFF processing temperatures. Intensities for -80 °C and -140 °C were shifted up by 50 and 100 counts respectively for readability.

Taken together, we speculate that the large total solid-air surface area and the more porous nature of the frozen thin films were responsible for the faster and more efficient removal of water from the frozen thin films, or in other words, the mass transfer from the frozen thin-films during freeze-drying was faster and more efficient in the primary drying phase. The mechanism of TFFD drying in the secondary drying phase is less understood because the secondary drying in conventional shelf freeze-drying is desorption dominated, which is usually considered heat-transfer driven (Pikal et al., 1990; Trelea et al., 2016). Because of the complexity of the sample stacking pattern, understanding the heat transfer mechanism is difficult. In general, heat is transferred to shelf frozen suspension mainly by conduction, with convection and radiation to some extents. However, heat is likely predominately transferred to the frozen thin films randomly arranged in a vial placed on the shelf of a freeze-dryer by radiation, with convection and conduction to a smaller extent. More experiments and modeling will have to be carried out to identify the extent to which every mode of heat transfer had contributed to the heat transfer during drying, especially in the secondary drying phase.

**Figure 8.**
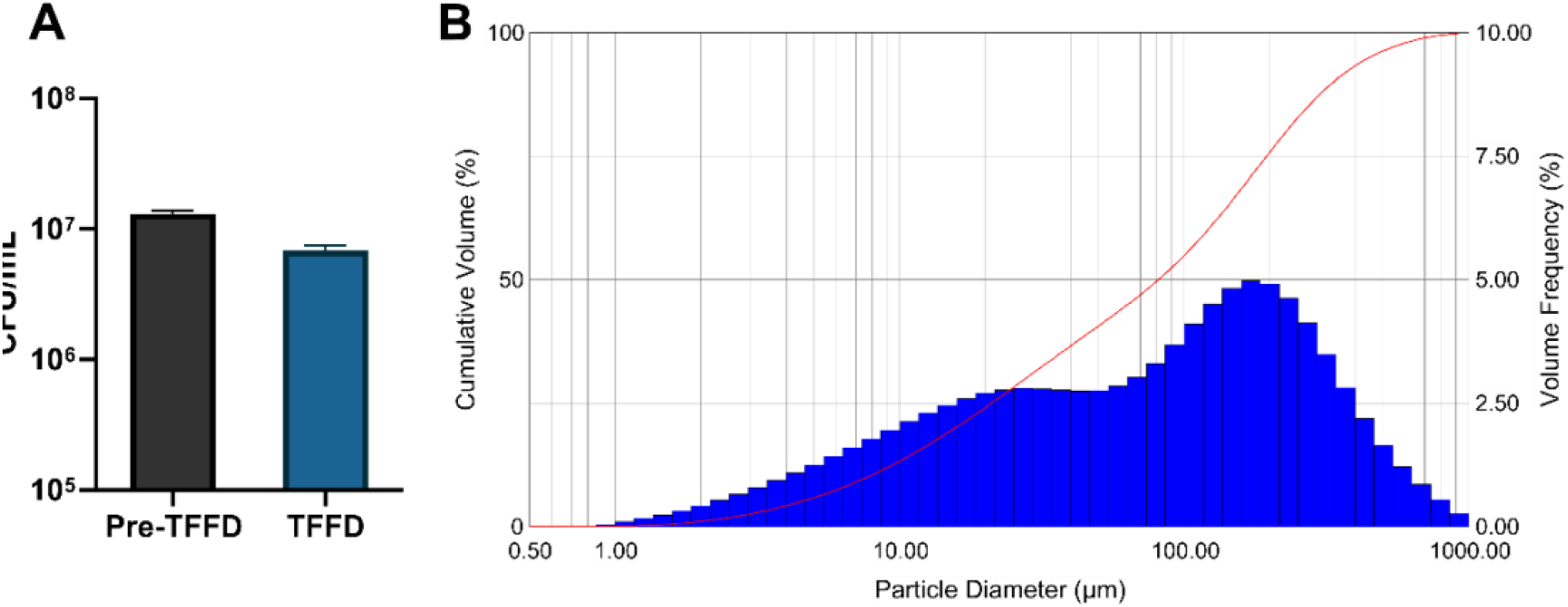
**(A)** *L. acidophilus* viability before and after being subjected to TFFD. (**B**) Particle size distribution of thin-film freeze-dried powder aerosol containing *L. acidophilus*.

Nevertheless, it is clear that at the 1 mL liquid in a 5 mL vial scale, water mass transfer from the frozen thin films during freeze-drying was significantly faster and more efficient than from shelf frozen suspensions, even with the small ice crystals generated by the ultra-rapid TFF process. On a larger scale, the speed and efficiency of water sublimation and desorption from frozen thin films, relative to conventional shelf frozen suspensions, will also likely be affected by how the frozen thin films are arranged in the container, the dimensions of the container (e.g., the diameter of the vials or bottles; vials or bottles vs. trays), and the heat diffusivity of the container. Also, as the films produced by TFF stack higher, the resistance to mass transfer between the films will reduce the drying rate, as shown in the previous experiment on freeze-drying pellets (Trelea et al., 2009). Therefore, there is still a potential limitation on container filling for the TFFD process.

### 3.4. Dry powder of *L. acidophilus* prepared by TFFD

Lastly, we tested the feasibility of using TFFD to prepare a dry powder of a bacterium other than *E. coli*. For this purpose, *L. acidophilus* was selected as it is Gram-positive and has various potential pharmaceutical and biomedical applications (Szatraj et al., 2017; Urbanska et al., 2007). To ensure the process parameters are transferable, we selected a sucrose-based excipient formulation with a composition similar to that used for *E. coli* (Fonseca et al., 2015). To maintain the bacterial viability while obtaining low moisture content, we used the thin-film freezing temperature of -40 °C (**Table 1**, #14) followed by a freeze-drying cycle with a short secondary drying time (**Table 2**, cycle 5). After the TFFD process, the log loss of bacterial viability was 0.28 (**Fig. 7a**). The relatively higher viability of the *L. acidophilus* after the TFFD process compared to *E. coli* was likely related to the fact that the Gram-positive lactobacillus bacteria have more rigid cell walls, leading to higher tolerance to the stress conditions in freezing and drying. The moisture content of the *L. acidophilus* TFFD powder was 1.18 ± 0.15% and the specific surface area was 4.692 m^2^/g (R^2^=0.999), both were comparable to *E. coli* TFFD powders.

Because *L. acidophilus* has been studied as a bacterial vector-based vaccine for potential mucosal immunization (Szatraj et al., 2017; Wang et al., 2020), we evaluated the particle size of the *L. acidophilus* TFFD powder after it was aerosolized using a dry powder sprayer for potential intranasal delivery. The particles demonstrated a bimodal distribution at around 30 µm and 200 µm (**Fig. 7b**). The Dv(10), Dv(50), Dv(90) values were 8.50 ± 1.6 µm, 92.3 ± 19.7 µm, and 283.5 ± 40.6 µm, respectively. Therefore, this powder formulation should have adequate aerosol performance properties for intranasal delivery as most of the particles were above 10 µm (Tiozzo Fasiolo et al., 2018; Xu et al., 2021, 2020).

## 4. Conclusion

In the present study, we showed that the TFFD technology can significantly increase the drying rate during freeze-drying. The main contributing factors include (i) the large total surface area of the thin films when randomly placed into a container or vial, promoting mass transfer through the void space between the films, (ii) the low thickness of the frozen thin films with lower resistance to mass transfer during freeze-drying, and (iii) the highly porous nature of the thin films allowing water molecules to easily escape during sublimation and desorption. In addition, we also demonstrated that the TFFD technology can be applied to prepare dry powders of bacteria suitable for potential intranasal administration while mitigating the bacterial viability loss within one log CFU/mL. Overall, TFFD technology is promising in addressing the drawback of time and energy inefficiency associated with conventional shelf freeze-drying.

## Acknowledgments

This work was supported in part by TFF Pharmaceuticals, Inc. and by the Mannino Fellowship in Pharmacy at UT Austin. We also would like to thank Dr. Zachary N. Warnken for his advice on powder aerosolization.

## Declaration of Interest

ROW and ZC report financial support provided by TFF Pharmaceuticals, Inc. ZC reports a relationship with TFF Pharmaceuticals, Inc. that includes: equity or stocks and funding. ROW reports a relationship with TFF Pharmaceuticals, Inc. that includes: consulting or advisory, equity or stocks, and funding. HX reports a relationship with TFF Pharmaceuticals, Inc. that includes: consulting or advisory. ZC, ROW, and JW have a patent pending to UT Austin.

